# RepeatParam: Algorithm for Parameterising Repeat Proteins and Analysis of Repeat Protein Architectures

**DOI:** 10.1101/2024.09.05.611252

**Authors:** Daniella Pretorius, James W. Murray

## Abstract

**Motivation:** Tandem repeat proteins consist of repetitive sequence and structure motifs and have diverse roles in Nature for molecular recognition and signalling. The architecture of repeat proteins can be described using simple helical parameters. Understanding these structural features can inform both the function of these proteins and be used to parametrically design new proteins. Despite their importance, no existing program is capable of completely parameterising repeat proteins.

**Results:** Here we describe a novel repeat protein parameterisation algorithm, RepeatParam, and a comprehensive repeat protein dataset. RepeatParam determines a helix that defines the global protein architecture and a superhelix that describes the relationship between consecutive repeats. We analyse the relationships between helical parameters and families of different repeat proteins.

**Availability:** https://github.com/dpretorius/repeat-protein-parameterisation

## Introduction

Tandem repeat proteins are very common in nature, constituting about 14% of all proteins [1], and are associated with over 50 human diseases [2]. These proteins are composed of 5-50 amino acid motifs that act as building blocks for a larger superstructure [3].

Repeat proteins can be categorised into 5 classes: crystalline aggregates, fibrous, elongated, closed and beads-on-a-string [4]. RepeatsDB, a database of tandem repeats in the Protein Data Bank (PDB), classifies proteins into these categories [4]. This paper focuses on elongated (α, αβ, and β-solenoids) and closed repeat proteins (**Figure 1a**). Unlike globular proteins, which fold into compact shapes, elongated repeat proteins form linear and extended structures, and closed repeat proteins form ring-like structures [5].

**Figure 1.**
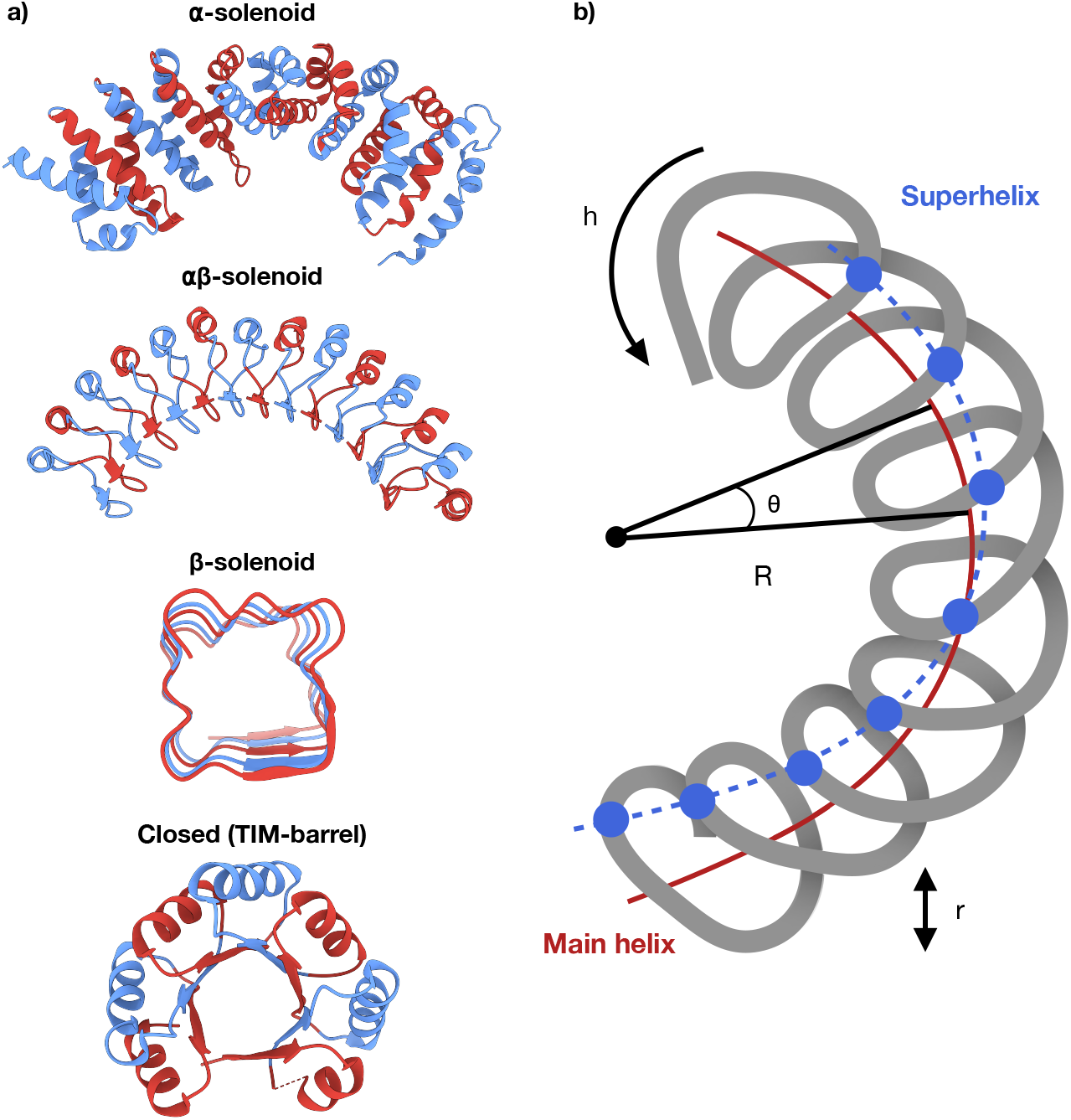
Repeat protein classes and architecture. **a)** Representative proteins for the repeat protein classes of interest: α-solenoid (PDB: 4UAD), αβ-solenoid (PDB: 4U06), β-solenoid (PDB: 2F3L) and closed repeats (PDB: 5BVL). In the case of the closed repeat a TIM-barrel is used as an example, but several other sub-families exist. Alternating repeats are shown in blue and red, non-repeat regions are shown in grey. **b)** Schematic representation of solenoid protein with helical parameters. The curvature of the protein can be described by the main helix (red). The superhelix (blue) connects equivalent positions in adjacent repeat units indicating the twist about the main helix. Repeat parameters notation: h = hand, R = radius, θ = twist and r = rise.

The structural diversity of solenoid and closed repeat proteins is significant. α-solenoid proteins are made up of repeated pairs of anti-parallel α-helices and have highly diverse shapes on a spectrum of curvature [6]. Key examples include armadillo, tetratricopeptide, HEAT, and ankyrin repeats [7]. α/β-solenoids combine α and β secondary structure elements, creating pronounced curvature due to the different widths of the α-helices and β-strands, as seen in Leucine Rich Repeats (LLRs) [8]. β-solenoid proteins are made from β-strands connected by tight β-turns. These structures typically have no or slight curvature as the distance between consecutive coils is determined by the set distance between β-strands of ∼4.8 Å [9]. The structural diversity of β-solenoid proteins is represented by the wide range of cross-sectional shapes differing in strand and turn composition [9]. Closed repeat proteins, such as triosephosphate isomerase (TIM) barrels and β-propellers, form compact ring structures. The number of repeats present for these folds is more constrained than open repeat proteins, with 4–10 repeats in a structure [10].

The modularity and symmetry of repeat proteins make them attractive targets for *de novo* design. Many previously designed proteins are tandem repeat proteins, with the majority being designed α-solenoids [11, 12, 13, 14]. Understanding the parameter ranges that define natural repeat proteins can guide the design of novel proteins with specific structural and functional attributes.

Understanding the structural characteristics of these proteins is crucial for both biological insights and applications in protein design. Elongated and closed repeat proteins can be described by the general definition set out by Kobe and Kajava (2000) for solenoid protein structures, where solenoid protein architecture can be modelled as a helix [7]. These proteins depend on the inter-unit interactions of tandem repeats to correctly fold. The repeat units follow a main helix that wraps about a central axis and the relative orientation of each repeat unit is described by a superhelix about the main helix (**Figure 1b**). Closed repeat proteins with toroidal shapes can be modelled as a helix with near-zero rise/pitch about the central axis. Using this generalised ‘helix-on-helix’ description, specific parameters are extracted to characterise the protein fold. For the main helix, these parameters include the radius (r), handedness (h), pitch (P), twist (t), and rise per repeat (s). For the superhelix, the parameters are omega (ω), the rotational frequency; alpha (α), the effective radius; and phase (ϕ), the angular offset. These parameters describe the overall protein fold [15] (**Figure 1b**). It should be noted that the nomenclature for helix and superhelix can vary in the literature, and we have chosen previously used convention [16].

Previous algorithms, such as HELFIT [17] and the Nievergelt method [18], have been developed for fitting points in 3D space to helical structures using total least squares methods. However, these general helix algorithms have not been applied to proteins and do not have the appropriate processing to parse a pdb format file, and do not consider superhelices. For proteins specifically, most efforts have targeted parameterisation at the level of individual α-helices, leaving a gap in methods for larger superstructures [19, 20].

Here we present a new algorithm, RepeatParam, that provides a parameterisation of repeat proteins and a dataset of repeat protein parameters from entries in the RepeatsDB database [4]. This dataset is useful for the general characterisation of repeat proteins and guiding parameter ranges for the future design of repeat proteins. Through this work, we aim to enhance the understanding of repeat protein architectures and their functional implications, facilitating advancements in protein engineering and design.

## Materials and methods

### Repeat database preparation

The classification and annotations from the RepeatsDB database were used for parameterisation [4]. RepeatsDB is a database of tandem repeat proteins from the PDB [21], categorised and annotated. Following a method similar to Brunette *et al*. (2015) [11], the residue ranges of repeats were extracted for α-solenoids (category 3.3), αβ-solenoids (category 3.2), β-solenoids (category 3.1), and closed repeats (category 4.1, 4.3, 4.4, 4.6 and 4.7).

A Python script was used to download PDB coordinates and extract the protein chain, start of repeat region, end of repeat region, average repeat unit length, index for each starting residue for repeat and index for insertion residues. These data were saved to a text file with command line arguments for automatic parameterisation. Proteins with fewer than four consecutive repeats or with a standard deviation greater than 3 in repeat unit lengths were omitted.

### Parameterisation algorithm

Repeat proteins were parameterised by fitting them to a ‘helix-on-helix’ model as outlined by Kobe and Kajava [7]. The main helix was fitted to the centroids of each repeat unit, and the superhelix was fitted to the starting Cα of each repeat (**Figure S1**).

RepeatParam processed PDB files with the following input arguments: filename, chain, start of repeat region, end of repeat region, average repeat unit length, index for each starting residue for repeat and index for insertion residues. This information was extracted from the RepeatsDB database. RepeatParam prepared structures by truncating non-repeat regions, removing heteroatoms and insertion regions, and normalising PDB residue indexing. The Cα atoms from the cleaned repeat regions were used to generate repeat unit centroids.

A general helix is described by 5 parameters: axis direction vector (ax, ay, az), radius (r) and pitch (P) (**Figure 2a**). These parameters for the main helix were obtained by fitting a helix to protein repeat unit centroids using a total least squares method adapted from the HELFIT algorithm by Enkhbayar *et al*. (2008) [17]. HELFIT, originally written for MATLAB, was adapted for RepeatParam using SciPy package in Python [22]. Refer to **Figure 2a** for notation and visualisation for fitting of the main helix.

**Figure 2.**
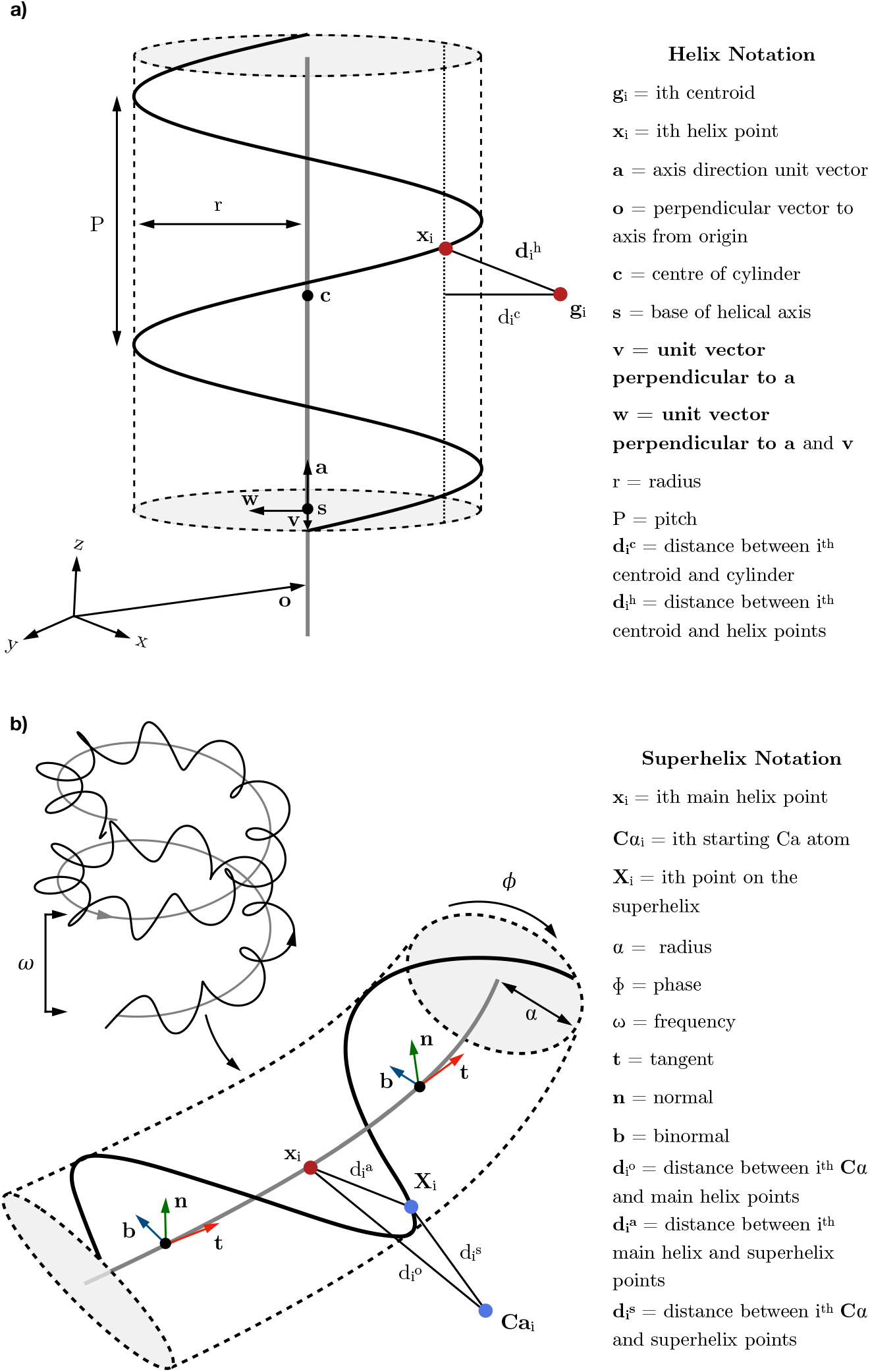
Main helix and superhelix fitting. **a)** (*Left*) Diagram of the fitting algorithm to centroid points to make the main helix. (*Right*) Associated mathematical notation. **b)** (*Left*) Diagram of the fitting algorithm to Cα atoms to make the superhelix. From the two rotations of main helix, a 90° section is focussed on. (*Right*) Associated mathematical notation.

The geometry of the superhelix is more complex than that of the main helix. The helical axis for the superhelix follows the main helix rather than a straight line. This curved helix can be described by the main helix path **x**(t) and 3 additional parameters: alpha (α), omega (ω) and phase (ϕ) (**Figure 2b**). Here, α is the effective radius, ω is the number of rotations per full rotation of **x**(t) and ϕ is the angular offset when t=0. A helix must be fitted to the starting Cα atom of every repeat to obtain these parameters for the protein superhelix. Refer to **Figure 2b** for notation and visualisation for fitting of the superhelix.

Details of the fitting methods for the helix and superhelix can be found in the **Supplementary Materials and Methods** section.

## Results and Discussion

### Parameterisation algorithm performance validation

RepeatParam determines a main helix that defines global protein architecture and a superhelix that describes the spatial relationship and orientation between consecutive repeats. To quantify the performance of RepeatParam, the Nievergelt dataset of 10 data points was processed and output parameters compared to existing algorithms. This dataset was previously used to assess the Nievergelt method and HELFIT. RepeatParam produced fits comparable to the previous methods (**Table S1**).

### Repeat protein parameter dataset

We generated a dataset of parameters for α-solenoids, αβ-solenoids, β-solenoids, and closed repeats. The quality of these parameters was assessed using the root-mean-squared deviation (RMSD) between the helix model and the empirical data, specifically for the centroids and the Cα atoms for the main helix and superhelix, respectively (**Figure 3a**). The mean RMSD for the main helix across all repeats was less than 2 Å. The mean RMSD for the superhelix fit is higher, at less than 5 Å across all repeat classes, reflecting the impact of errors propagated from the main helix fit and the inherent irregularities of the Cα atoms forming the superhelix. Additionally, an uncertainty metric was calculated for each parameter to further assess the quality of the fits.

**Figure 3.**
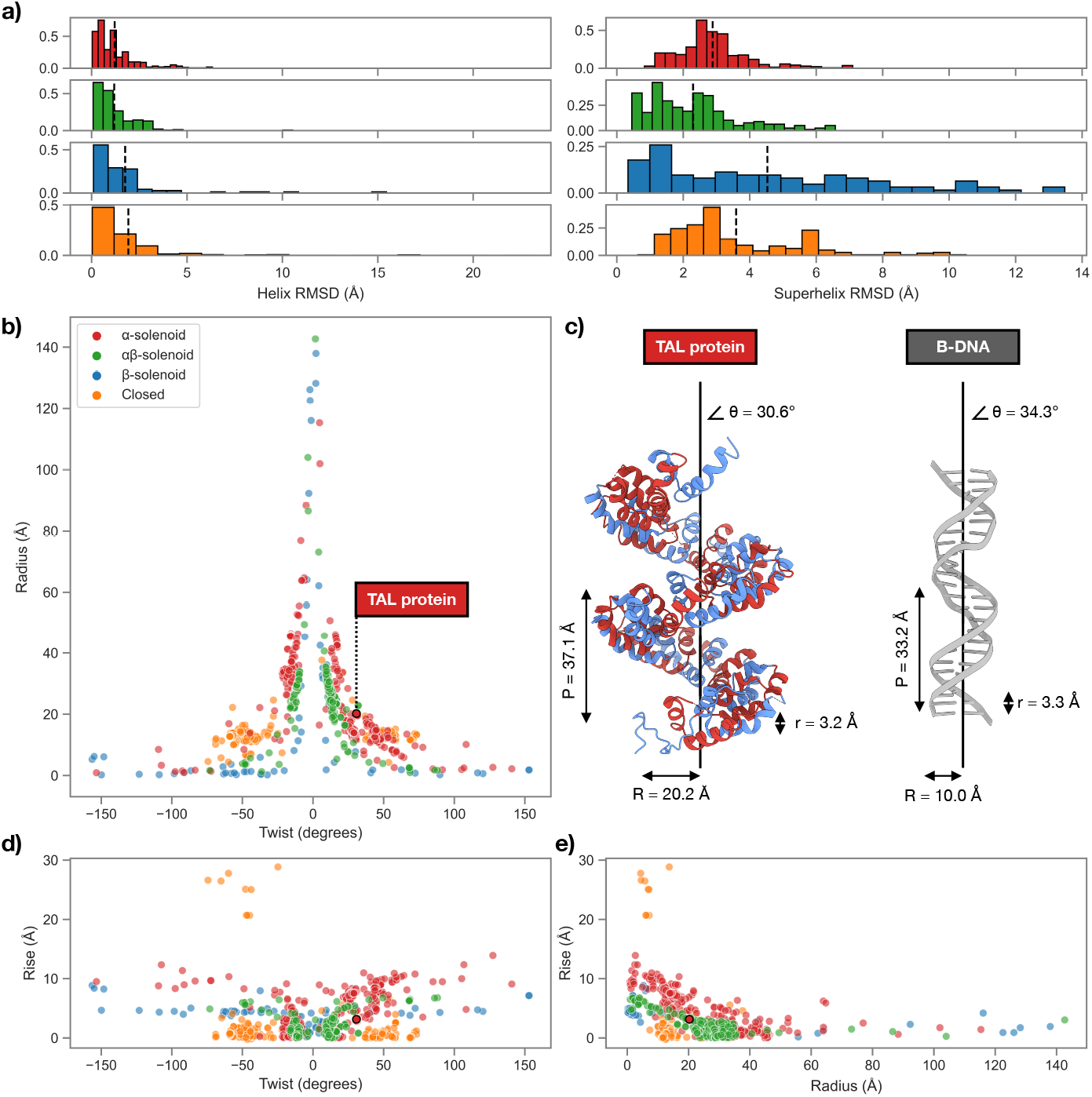
Tandem repeat protein parameter dataset analysis. **a)** RMSD histogram for main and superhelix model. The comparative RMSD fitting of the main and superhelix compared to the data for each repeat protein class. Mean RMSD is shown by the black dotted lines. The data was filtered by excluding any fitting > 2 standard deviations from the mean. Bin size of 20 for all histograms. **b)** Distribution between twist and radius, DNA binding TAL-protein highlighted. **c)** Comparison of helical parameters: (*Left*) for the DNA-binding TAL effector protein (PDB: 3UGM); (*Right*) for the corresponding B-DNA in the same crystal structure. **d)** Distribution between rise per repeat and twist. **e)** Distribution between radius and rise. For the data in plots **b, d, e)** only examples where the model was a good fit, helix RMSD > 2 Å was used.

Parameterising repeat proteins with significant irregular regions proved more challenging. The β-solenoid class, which had high irregularity, exhibited the highest RMSD among all repeat proteins, indicating a poorer fit. This class is also underrepresented in our dataset, with fewer examples compared to other repeat types. However, with the recent expansion of protein structure databases with the AlphaFold Database [23] and improvements in tandem repeat protein identification methods with machine learning [24, 25], there will be an expansion of the number of well-characterised repeat proteins, providing a larger pool of candidates for analysis.

For all further analysis, we filtered the parameter dataset to include only those with an average RMSD of less than 2 Å to the main helix.

### Repeat protein parameter relationships

The independent parameters for the main helix include pitch (P) and radius (r). For the superhelix, the independent parameters are effective radius (α) and rotational frequency (ω). From the main helix parameters, derived parameters such as twist (the angular displacement per repeat) and rise per repeat (the linear displacement along the helix axis per repeat) were calculated. The relationships between the main helix parameters or radius, twist and rise were plotted (**Figure 3b, d, e**).

Most superhelices exhibit a very small rotational frequency, due to minimal twist between consecutive repeats. However, there are exceptions; for example, in the armadillo repeat protein (PDB: 5EKG), the superhelix completes a quarter turn for each full turn of the main helix (**Figure S2**).

The pairwise repeat parameter comparisons in **Figure 3** highlight the physical limitations of repeat proteins and the differences between repeat classes. Brunette *et al*. (2015) suggested that the helical parameters of radius (R), twist (θ) and rise per repeat (r) are correlated [11]. The total length of a repeating unit can be approximated by:

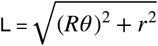

This approximation applies to helical proteins with a non-negligible rise, so closed repeat proteins do not follow many of the expected trends. Assuming that the total length of a repeat unit remains constant while global parameters change, proteins with high twist tend to have a smaller radius. The inverse relationship between twist and radius is shown in **Figure 3b**. No overall trend is represented between twist and rise (**Figure 3d**), but many β-solenoid proteins cluster in a band at a rise of ∼4-5Å, which is the approximate distance between parallel β-strands in a sheet (4.9 Å) [26]. It is clear from both plots that zero twist is very rare in natural proteins. Also the left-handed (negative twist) and right-handed (positive twist) data mirror each other, displaying comparable parameter values irrespective of handedness. Similar distributions of radius and rise compared to twist were seen for α-solenoids in previous work [27].

The relationship described above for the total length of a repeating unit is generally seen between radius and rise, where proteins with a larger radius tend to have smaller rise (**Figure 3e**). Closed repeat proteins deviate from this trend, having a small rise irrespective of radius due to their circular geometry. These closed repeats fill an architectural gap not accessible to other repeat protein types.

In a DNA-binding α-solenoid TAL effector protein (PDB: 3UGM) [28], the parameters from RepeatParam closely match those of B-DNA [29] (**Figure 3c**). For the TAL effector, the twist (30.7°) is comparable to the rotation of B-DNA per base pair (34.3°). The rise per repeat (3.2 Å) is consistent with B-DNA’s rise per base pair (3.3 Å). The larger radius for the protein (20.2 Å) to B-DNA’s radius (10 Å) is due to the repeat centroids being in the middle of the repeat units, providing space for the protein to fit around the DNA structure. This compatibility with the DNA groove is critical for the protein’s function, as each repeat in the TAL protein identifies a single base of DNA. Understanding these parameters is essential for comprehending the biological function of solenoid proteins and provides guidelines for designing proteins with specific functions, such as DNA binding.

### Repeat protein families

A t-SNE projection of the parameters that describe the repeat protein helices was used to reduce the dimensionality of the repeat parameter dataset and identify patterns for the 4 variables describing the main helix (**Figure 4**). α-solenoids show the highest variation as they span many geometries [6], examples highlighted to show this diversity are the large radius tetratricopeptide (PDB: 1O1R) which requires a large interaction surface to bind to large protein substrate [30], and the tetratricopeptide (PDB: 5MFH) with a small radius but large rise per repeats for helix binding at the concave and convex surfaces [31, 32]. Closed repeats cluster in two distinct groups, the parameters shared between all closed repeats are very similar, however the twist between repeats can differ significantly, in the case of two different β-propeller proteins with seven repeat (PDB:6OSA) and six repeat (PDB:5H47) in different clusters due to their different twists [33, 34]. Most αβ-solenoids occupy a similar position in parameter space as LRR are the dominant αβ-type, which have large radius and curvature, as seen by the two LRR examples (PDB: 4BSS, 3TSR) [35, 36]. As the β-solenoid parameterisation was the poorest and there were significantly less examples used, the clustering is less obvious. However, most of the β-solenoid proteins cluster in a band together with some low curvature αβ-solenoids, all of these proteins have a very small radius, and have a consistent rise of ∼4-5Å, as seen by an R-type (PDB: 2G0Y) and T-type (2VHE) β-solenoids [37, 38].

**Figure 4.**
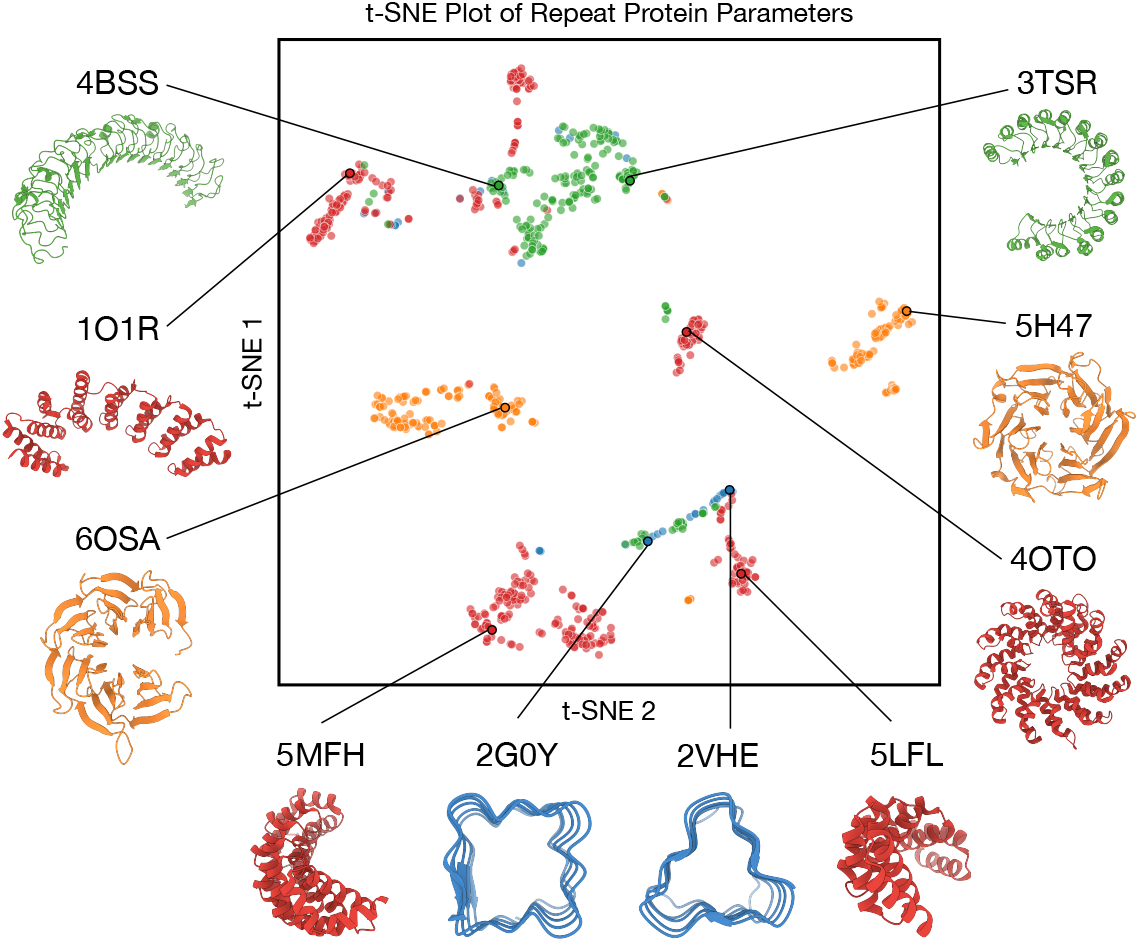
Repeat protein family distribution split by helical parameters. t-SNE projection of repeat helical parameters for α-solenoids (red), αβ-solenoids (green), β-solenoids (blue), and closed (yellow) repeats. PDB IDs of each protein are displayed above each structure.

## Conclusion

RepeatParam is an algorithm for parameterising repeat proteins, addressing a gap in bioinformatic tools by defining protein architecture through main helix and superhelix parameters. With RepeatParam we generated a comprehensive dataset of repeat protein parameters, serving as a valuable resource for future work in the repeat protein area. RepeatParam’s ability to handle diverse repeat classes and integrate with emerging tools makes it a critical resource for understanding repeat protein architectures and further usage for repeat protein design applications.

## Notes

### Competing Interest Statement

The authors have declared no competing interest.

