## Supplementary material for "RepeatParam: Algorithm for Parameterising Repeat Proteins and Analysis of Repeat Protein Architectures": Figure S1, Figure S2, Table S1, Supplementary Materials and Methods

**This PDF includes:**

- Materials and Methods
- Supplementary table (S1)
- Supplementary figures (S1 & S2)

### Materials and methods

#### Main helix fitting

Refer to **Figure 2a** through this section for notation and visualisation for fitting of the main helix.

Stage 1 - Initial axis and radius determination. It is critical that the initial axis direction vector (**a**) estimate is good to avoid being trapped in a local minimum while fitting. The initial axis direction is calculated using the Khan method [1]. This takes the bisector of the angle between three consecutive helical points to create a perpendicular vector to the estimated helical axis. Four points are required to yield two vectors perpendicular to the helix axis, the cross product of these vectors gives the helix axis. Thus, the minimum number of repeating units in a target protein is 4. A projection matrix of the initial axis direction vector was then used to determine the centre (**c**) and radius (r) of a right circular cylinder that approximates the helix [2].

Stage 2 - Cylinder fitting. The perpendicular vector from the origin to the initial axis (**o**) is found using by projecting the origin (O) onto the axis:

$$\mathbf{o} = \mathbf{c} + (O - \mathbf{c}) \cdot \mathbf{a})\mathbf{a} \tag{1}$$

The orthogonal vector from the axis to the $i^{th}$ centroid point ($\mathbf{r}_i$) is found using the same method as (1) by projecting the centroid point onto the axis:

$$\mathbf{r}_i = \mathbf{g}_i - \mathbf{o} - (\mathbf{g}_i \cdot \mathbf{a})\mathbf{a} \tag{2}$$

The direction of $\mathbf{r}_i$ represents the orthogonal distance from the axis to the $i^{th}$ centroid point. The distance from the $i^{th}$ centroid point to the surface of the cylinder ($d_i^c$) is given by the difference between the distance to the axis and the cylinder radius:

$$d_i^c = |\mathbf{r}_i| - r \tag{3}$$

The cost function ($J^c$) is the sum of squares of the distance to the cylindrical surface:

$$\mathbf{J}^c(r, \mathbf{a}, \mathbf{o}) = \Sigma(d_i^c)^2 \tag{4}$$

This is subject to the constraints that vectors **a** and **o** are perpendicular to each other and that **a** is a unit vector

Stage 3 - Pitch and handedness determination. The centroids are translated so that **c** is at the origin. To simplify calculations, the centroids were rotated with a rotation matrix to align **a** with the z-axis. The relative displacement (z) along the z-axis and the angular rotation (θ) around the z-axis are calculated for each centroid. This is done by projecting the centroids onto the z-axis for displacement and onto the x-y plane for rotation. The direction of the angular rotation (clockwise or counterclockwise) indicated the right or left-handedness of the protein. Displacement and rotation are described using a simple linear regression model:

$$z = m\theta \tag{5}$$

A quadratic loss function measuring the error in the model is minimised using a least squares method for the gradient (m):

$$m = \frac{\Sigma(\theta_i - \bar{\theta})(z_i - \bar{z})}{\Sigma(\theta_i - \bar{\theta})^2} \tag{6}$$

Using m, the pitch (P) of the helix is determined for a full rotation (θ=2π):

$$P = 2\pi m \tag{7}$$

Handedness is given by the sign of the pitch. For RepeatParam all pitch values are taken as positive for further fitting. If the estimated pitch is determined to be negative, then it is made positive by using the opposite sign of **a** and changing the handedness accordingly.

Stage 4 - Full minimisation with all parameters. The rotation around the helix axis (t) for the centroid points is determined by parameters that are minimised during optimisation. This approach allowed t to be minimised without increasing the number of variables in the optimisation. The vector to each centroid point from helical axis ($\mathbf{u}_i$) is found in the same way as (2):

$$\mathbf{u}_i = \mathbf{g}_i - \mathbf{o} - (\mathbf{g}_i \cdot \mathbf{a})\mathbf{a} \tag{8}$$

The angles between pairwise vectors $\mathbf{u}_i$ and $\mathbf{u}_{i+1}$ is calculated:

$$t_i = \cos^{-1}\left(\frac{\mathbf{u}_i \cdot \mathbf{u}_{i+1}}{||\mathbf{u}_i||\,||\mathbf{u}_{i+1}||}\right) \tag{9}$$

The starting point of the helix (**s**) is defined by the projection of the first data point ($\mathbf{t}_0$) onto the helix axis. The first centroid point is used an approximation of the first helix point:

$$\mathbf{s} = \mathbf{o} + ((\mathbf{t}_0 - \mathbf{o}) \cdot \mathbf{a})\mathbf{a} \tag{10}$$

**v** is the unit vector defining the direction from **s** to $\mathbf{t}_0$. **w** is perpendicular to both **a** and **v**, calculated as their cross product. These three vectors form the orthogonal system that the helix was built from. The conditions are always true that:

$$\mathbf{a} \cdot \mathbf{v} = 0 \qquad \mathbf{v} \cdot \mathbf{w} = 0; \ |\mathbf{v}| = 1 \qquad \mathbf{w} \cdot \mathbf{a} = 0; \ |\mathbf{w}| = 1 \tag{11}$$

Using these values, the main helix (**x**(t)) is described by:

$$\mathbf{x}(t) = \mathbf{s} + \frac{\mathbf{a}Pt}{2\pi} + r(\mathbf{v}\cos(t) + \mathbf{w}\sin(th)) \tag{12}$$

For right-handed proteins, h=1; otherwise, h=−1.The orthogonal distance between helix and centroid points ($d_i^h$) is given by:

$$d_i^h = |\mathbf{g}_i - \mathbf{x}_i| \tag{13}$$

The cost function ($J^h$) for the full minimisation was defined as the sum of squares of the distances between the main helix (model) and centroids (data). This function was minimised for the 11 parameters P, r, **a**, **o**, $\mathbf{t}_0$ subject to the constrains (5) and (6):

$$\mathbf{J}^h(P, r, \mathbf{a}, \mathbf{o}, \mathbf{t}_0) = \Sigma(d_i^h)^2 \tag{14}$$

**Superhelix fitting**

Refer to **Figure 2b** through this section for notation and visualisation for fitting of the superhelix.

**Stage 1 - Project Ca coordinates onto main helix.** The starting Cα points are projected onto the main helix since the superhelix is built around it:

$$d_i^o = |\mathbf{x}_i - Ca_i| \tag{15}$$

This orthogonal distance is found by determining the parameter n that minimises the distance between the $i^{th}$ Cα and main helix curve. The parameter n represents the position along the main helix path x(t) where the Cα coordinates are projected. The cost function ($J^h$) is minimised to find the optimal n that gives the smallest distance between the Cα points and the helix:

$$\mathbf{J}^h(\mathbf{n}) = \Sigma(d_i^o)^2 \tag{16}$$

Using these optimal values of n, a new set of points on the main helix corresponding to the starting Cαs (T) is determined:

$$\begin{aligned} \mathbf{n} &= \{n_1, n_2, n_3 \dots n_N\} \\ \text{therefore} \quad T &= \mathbf{x}(n) \end{aligned} \tag{17}$$

**Stage 2 - Frenet frame on the main helix**. A Frenet frame is an ordered orthonormal triple of vectors (**t, n, b**) that describe a new coordinate system relative to any point on a curve [3]. This orthogonal system serves the same purpose as the (**a**, **v**, **w**) vector triple that build a general helix. The Frenet frames for each projected point on the main helix **x**(n) are defined by:

$$\mathbf{t}(n) = \frac{\mathbf{x}'(n)}{||\mathbf{x}'(n)||} \qquad \mathbf{n}(n) = \frac{\mathbf{t}'(n)}{||\mathbf{t}'(n)||} \qquad \mathbf{b}(n) = \mathbf{t}(n) \times \mathbf{n}(n) \tag{18}$$

Stage 3 - Radius and phase determination. The distance between each $\mathbf{x}_i$ and $C\alpha_i$ ($d_i^a$) is given by:

$$d_i^a = |Ca_i - T_i| \tag{19}$$

The average $d_i^a$ over the N Cα atoms is used to find the effective radius (α)

$$\alpha = \frac{\Sigma(d_i^a)}{N} \tag{20}$$

To find the phase, the estimated position of the first superhelix point with 0 phase (Q) is determined. The first main helix point ($T_1$) and Cα atom ($Ca_1$) are represented as A and P respectively:

$$Q = A + \cos(0)\mathbf{n} + \sin(0)\mathbf{b} \tag{21}$$

The normal vector (N) to the plane that A and P lie on is found taken the cross product:

$$N = (\overrightarrow{AP} \times \overrightarrow{AQ})m \tag{22}$$

N must be the same general direction as the tangent vector (**t**) of A. Thus, if N·t < 0 then m=-1, otherwise for N·t > 0, m=1. The angle ∠PAQ (ϕ) is determined in the same way as in (9):

$$\phi = \cos^{-1}(\frac{\overrightarrow{AP} \cdot \overrightarrow{AQ}}{||\overrightarrow{AP}|| \, ||\overrightarrow{AQ}||}) \tag{23}$$

ϕ must be taken clockwise relative to the direction of the main helix. To do this, a reference point of the normal vector, which faces into the main helix, is used. The vectors AP, AQ, and N are stacked into a 3x3 matrix (M). If det(M) < 0 then ϕ can be used, otherwise the corresponding angle is taken:

$$\begin{aligned} \phi &= \phi, && \text{if det}(M) < 0 \\ \phi &= 2\pi - \phi, && \text{if det}(M) > 0 \end{aligned}$$

Stage 4 - Full minimisation with all parameters. The superhelix (X(n,θ)) can be described by:

$$X(n, \theta) = \mathbf{x}(n) + \alpha(\cos(\theta)\mathbf{n}(n) + \sin(\theta)\mathbf{b}(n)) \tag{24}$$

$$\theta = n\omega + \phi \tag{25}$$

The orthogonal distance between superhelix and starting Cα points ($d_i^s$) is given by:

$$d_i^s = |Ca_i - X_i| \tag{26}$$

The cost function ($J^s$) is the sum of squares of the distances between the superhelix (model) and starting Cα atoms (data). This is minimised for the three parameters α, ω and ϕ, with the constraints 0 < α and 0 < ω < 2π:

$$\mathbf{J}^s(\alpha, \omega, \phi) = \Sigma(d_i^s)^2 \tag{27}$$

### Supplementary Table

**Table S1 | Comparison of helix fitting parameters on the Nievergelt dataset.**

| Objective | RepeatParam | HELFIT | Nievergelt |
|---|---|---|---|
| Direction vector (**a**) | (0.32, 0.72 0.62) | (0.32, 0.73, 0.61) | (0.33, 0.67, 0.67) |
| Helix radius (r) | 181 | 178 | 195 |
| Helix RMSD (Å) | 0.68 | 0.59 | 0.72 |

The Nievergelt data lie on a cylinder surface with a radius of 195 AU (arbitrary units) and axis direction parallel to (0.33, 0.67, 0.67). This assesses the main helix fitting method.

### Supplementary Figures

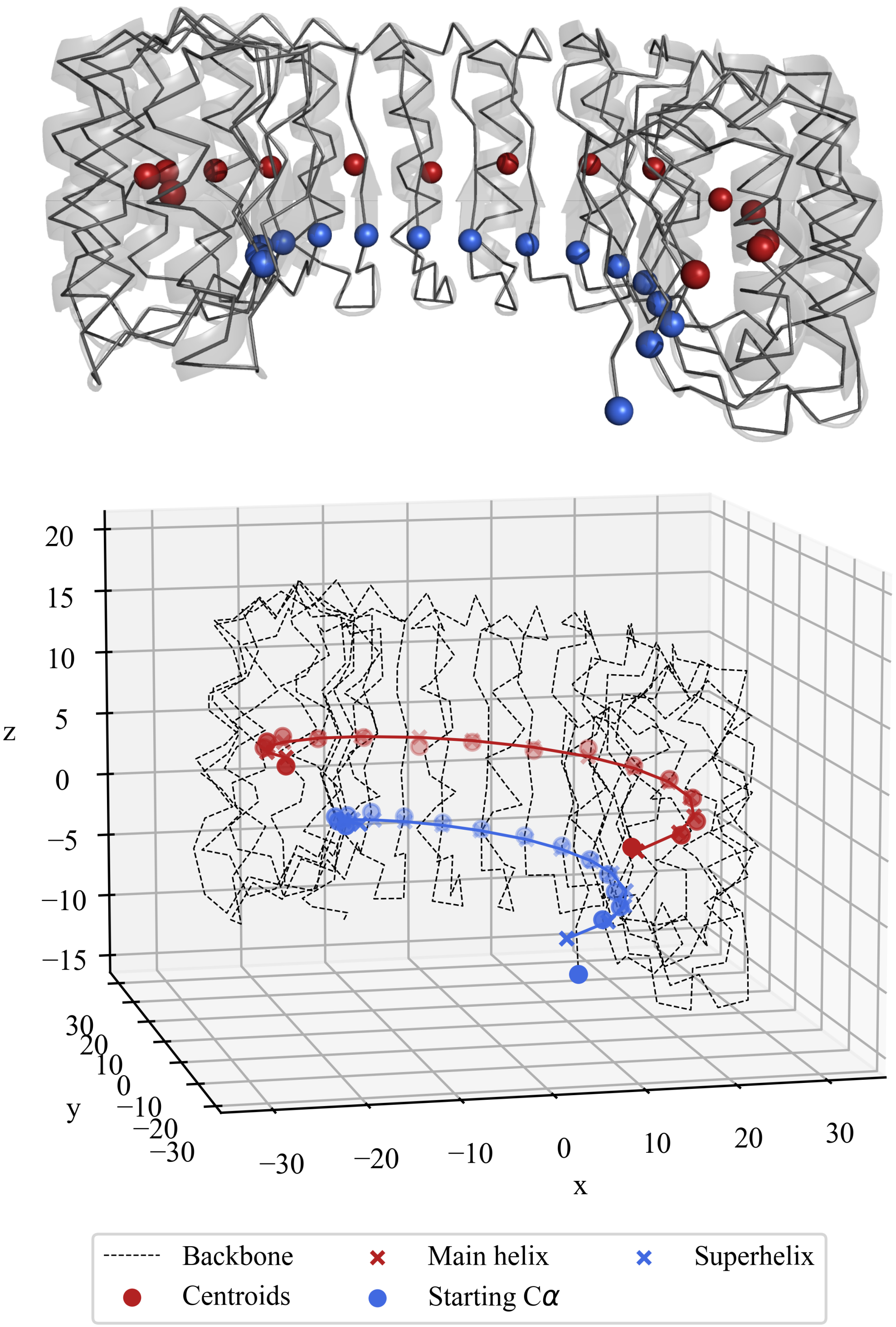


**Figure S1 | Parameterisation algorithm.** (*Above*) Pymol representation of an αβ-solenoid shown with spheres for the centroids of each repeat (red) and the starting Cα for each repeat (blue). (*Below*) The same αβ-solenoid shown as parameterised by my algorithm. The main helix is represented in red and the superhelix is represented in blue. The backbone trace of the Cα atoms are shown by the dotted line. (PDB: 2BNH)

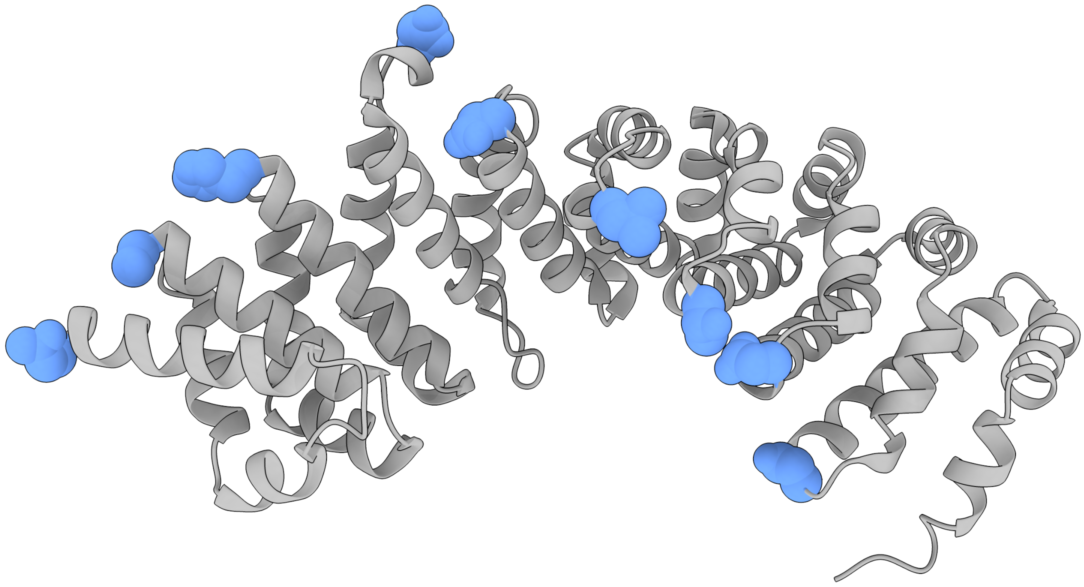

**Figure S2 | Superhelix path of armadillo repeat protein.** Pymol representation of armadillo α-solenoid shown with spheres for the starting Cα for each repeat (blue), this shows the path of the superhelix, which completes a quaternary turn for each full turn of the main helix. (PDB: 5EKG)
